# Mechanism of auxin-dependent gene regulation through composite auxin response elements

**DOI:** 10.1101/2024.07.16.603724

**Authors:** Daria D. Novikova, Nadya Omelyanchuk, Anastasiia Korosteleva, Catherine Albrecht, Viktoriya V. Lavrekha, Dolf Weijers, Victoria Mironova

## Abstract

The plant signaling molecule auxin controls growth and development, largely through activating and repressing the expression of thousands of genes. Auxin-dependent transcriptional changes are mediated by DNA-binding Auxin Response Factors (ARF), whose AuxRE DNA binding sites are well-known. The identification of the first AuxRE showed this to be part of a composite element with a second motif. Indeed, systematic analysis showed other DNA motifs to be enriched in auxin-regulated promoters. Neither the basis for this enrichment nor the mechanisms for the activity of composite AuxRE’s is known. Here, we systematically mined Arabidopsis promoters for composite AuxRE elements enriched in auxin-responsive genes. We identified many and show that their presence is a reliable predictor of auxin response. Through mutating these elements and their higher-order modules, we demonstrate function in promoter activity. Lastly, we identified transcription factors (TFs) that bind AuxRE-associated motifs, showed their involvement in auxin response, and discovered that several of these TFs directly bind ARF proteins. We propose that ARF-TF complexes specifically bind compound motifs in promoters, and act as a source of diversification in auxin-dependent gene regulation.

## Introduction

The plant signaling molecule auxin controls essentially all aspects of growth and development, and developmental contexts determine its many unique responses^1^. The core mechanism of transcriptional response to auxin starts with the binding of auxin to its receptors from the TRANSPORT INHIBITOR RESPONSE1/AUXIN SIGNALING F-BOX (TIR1/AFB) family, which facilitates the ubiquitination and degradation of AUXIN/INDOLE-3-ACETIC ACID (Aux/IAA) transcriptional inhibitors^2,3^. Aux/IAAs do not bind DNA on their own but dimerize with the AUXIN RESPONSE FACTOR (ARF) transcription factor (TF) family^4^. Upon an increase in auxin concentration, ARFs that are already anchored to their binding sites (AuxRE; Auxin Responsive Elements) are released from AUX/IAA-mediated inhibition and can regulate transcription of their targets^5,6^.

Although plants can have large families of ARFs and Aux/IAAs, it is unclear if this is enough to account for the complexity and diversity of processes that auxin regulates^1,7^. Clearly, combinatorial interactions of auxin signal transduction components can be created by (1) tissue- and organ-specific expression of different auxin receptors, ARFs, and Aux/IAAs^8,9^, (2) half-lives of proteins and their auxin-sensitivity^10,11^, (3) affinities of proteins to each other^9^, (4) affinities of the TIR/AFB receptors to auxin, and (5) interaction with other protein families^12,13^, including interactions with other TFs^14^. All these can contribute to the assembly of functionally distinct response machineries in cells. Likewise, variation in DNA elements can contribute to diversity in auxin responses. ARFs bind a TGTCNN motif, with TGTCGG displaying the highest affinity^15–17^. ARFs form symmetric dimers with their DNA-binding Domains (DBD), and this explains the cooperative binding to a precisely spaced, inverted repeat of AuxRE’s^17^. More generally, the binding affinity, ARF-specificity, and auxin response strength greatly depend on the syntax (spacing and orientation) of two ARF-binding elements^16–19^.

A view that integrates protein-centered and DNA-centered perspectives would propose the recruitment of ARFs together with non-ARF TFs to composite cis-regulatory elements having respective binding sites located in close proximity^20,21^. There are a number of examples of such ARF/non-ARF interactions on DNA that mediate tissue-specific auxin responses. For instance, petal growth is regulated by ARF8 and the bHLH TF BIGPETALp^22^; gynoecium development is promoted via ARF3 together with another bHLH factor, IND^23^, and fruit valve growth depends on interactions between ARF8 and ARF6 with the MADS transcription factor FUL^24^. Auxin and brassinosteroid responses converge via ARFs and the BES1/BZR1 TFs^25–27^. GBF factors from the bZIP family bind to ARF5 to modulate expression of vascular genes^28^.

Such ARF-TF interactions should reflect in the arrangement of DNA motifs in the promoters of auxin-regulated genes, as both would have to bind in close vicinity to cooperatively regulate the targets. Indeed, ARF and GBF binding sites are found in close proximity^28^, but it is unclear if this is a general mechanism underlying auxin-dependent gene expression. AuxRE’s were first identified through detailed *GH3* promoter analysis in soybean^29,30^. This early work showed that the minimal AuxRE could not mediate expression without the presence of a second (coupling) element^30^. Here, the second element (CACG) provided constitutive activity, which was rendered auxin-responsive through the presence of the AuxRE. While the AuxRE was later shown to bind ARFs^15^, the TF that binds and regulates the coupling element was not found. Given that for most cases where ARF-TF interaction have been shown, it is unclear if these are relevant for the control of shared targets through composite elements, it is largely an open question if these mechanisms contribute to diversity in auxin response.

Here, we explore this hypothesis. We mined the Arabidopsis genome for composite elements (AuxRE-motifX) enriched in auxin-responsive promoters. We functionally validate a number of these, identify TF’s binding the coupling elements and demonstrate that several of these are relevant for auxin response. Lastly, we find that several TFs bind directly to ARFs, suggesting ARF-TF complex binding to composite DNA elements as a widespread mechanism underlying auxin-dependent gene regulation.

## Results

### Systematic identification of composite auxin-responsive elements

We assessed the upstream regions of auxin-activated and auxin-inhibited genes taken from 19 transcriptomic experiments on auxin treatments of Arabidopsis seedlings (Supplementary Table 1; Fig. 1a). The datasets were grouped into early (<=2 hours) and late (>2 hours) responses and analyzed separately. With *metaRE*^31^, we searched for composite (henceforth “bipartite”) AuxRE’s (biAuxRE’s) that were systematically (meta-p-value < 1E-5) and significantly (permut-p-value < 0.05) overrepresented in auxin early- and late-responsive promoters for up- and down-regulated genes. The biAuxRE’s had the following structures: TGTCNN-spacer-NNNNNN or NNNNNN-spacer-TGTCNN, where N could be any nucleotide (A, T, G, C), and the spacer length between the ARF-binding core and the coupling motif was set from 0 to 15 bp. In total, 31200 possible biAuxRE were assessed, and 591 of them were found to be enriched in the promoters of auxin-regulated genes, 398 in activated, 191 in repressed, and two in both (Supplementary Table 2). More than 80% of the enriched biAuxRE’s were found in promoters of early auxin-responsive genes (347 in early-activated and 131 in early-inhibited) (Fig. 1b), and only 13 biAuxRE’s belonged to both early and late response categories. This aligns well with the role of ARFs in the primary response to auxin^5^.

**Figure 1.**
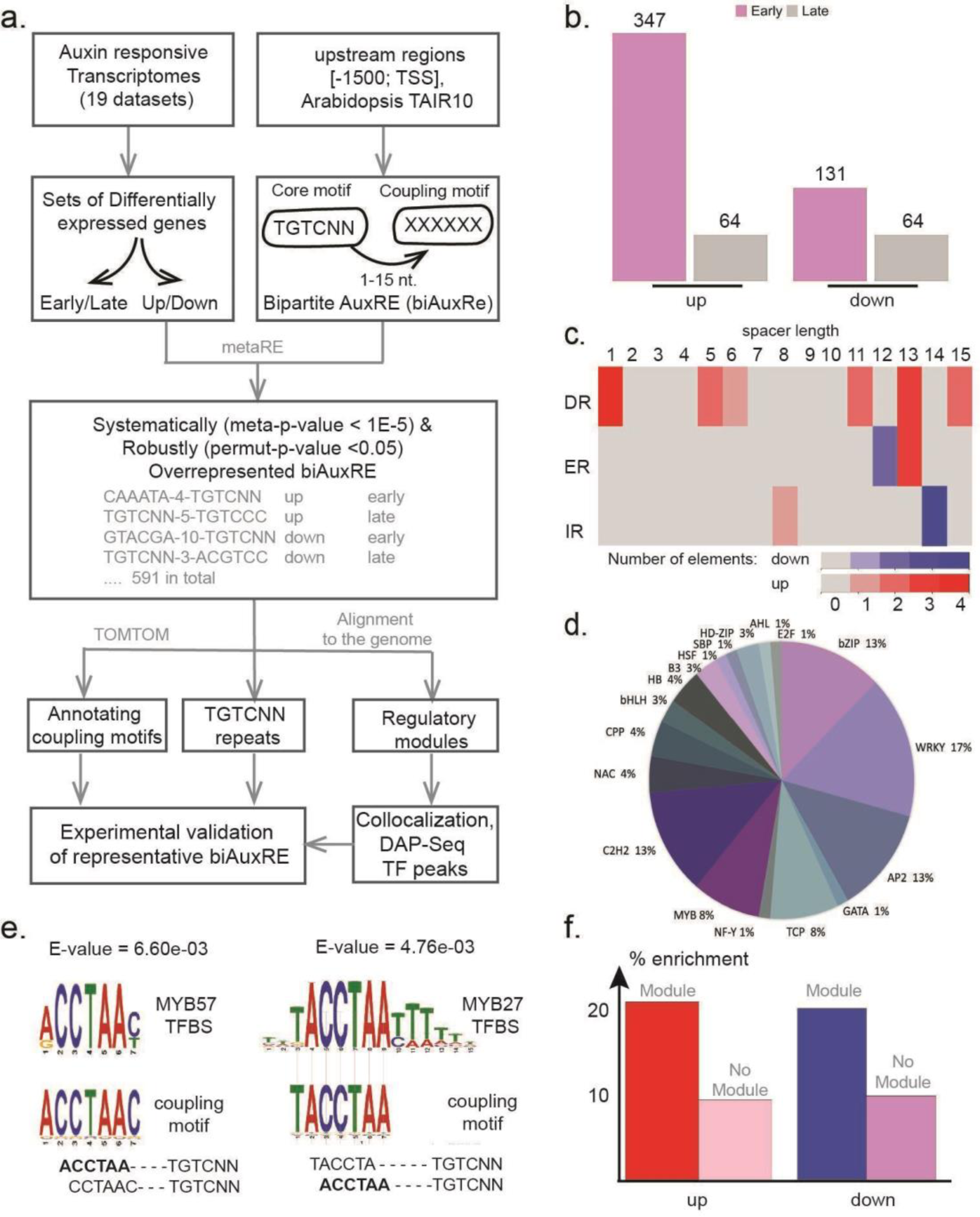
Identification of an auxin-responsive regulatory code. **a**. Graphical summary of the bioinformatic pipeline. **b**. The number of biAuxRE associated with up/down and early/late responses to auxin in the analysis with *metaRE*^31^. **c**. Overrepresented TGTC-containing repeats, total number of repeats is highlighted by colour. DR: direct repeat; ER: everted repeat; IR: inverted repeat. **d**. Coupling motifs annotation performed with the TOMTOM^33^. The annotated motifs are grouped by the TF family for which binding sites a significant resemblance (E-value <0.05) has been detected. **e**. An example of two distinct colocalizations of biAuxRE’s in gene promoters with a shift in the coupling motif position. The extended motifs significantly match different MYB-binding sites (TOMTOM, E-value <0.05). **f**. The genes with the predicted regulatory modules in their upstream regions are two-times more likely to respond to auxin with either activation or inhibition than the genes without the predicted module.

We next validated the inference by exploring if the analysis detected AuxRE repeats that are known to mediate auxin response. Indeed, there were 25 bipartite elements containing two TGTCNN sequences, which differ in orientation, spacer, and the sequence of the second motif (we allowed a shift in the coupling motif: **TGTC**NN, N**TGTC**N, or NN**TGTC**) (Supplementary Table 3). The identified elements are grouped into 10 types of repeats: direct repeats (DR) with spacers 1, 5-6, 11, 13, 15; inverted repeats (IR) with spacers 8 and 14, and everted repeats (ER) with spacers 12-13 (Fig. 1c). Notably, we detected all the repeats that have been shown earlier to control the auxin response: DR5, IR8, and ER13^16,19,32^. The most abundant repeat associated with auxin-dependent gene activation was a yet unknown DR1 with four biAuxRE’s of this type detected in total. Interestingly, we also detected two elements specifically associated with down-regulation, ER12 and IR14, neither of which has been described before.

Next, we annotated the coupling elements from each biAuxRE with the TOMTOM tool^33^. 12% of the identified biAuxRE were found to contain coupling elements significantly matching known TF binding sites (*E*-value <0.05) (Supplementary Table 4). The most numerous were the motifs for the WRKY, AP2/ERF, bZIP, C2H2, MYB, TCP, and HD-ZIP families (Fig. 1d).

As TF binding site models are usually longer than 6 nucleotides, we hypothesized that some of the coupling elements might be fragments of more extended cis-elements. Indeed, the alignment of biAuxRE’s relative to each other revealed longer coupling motif variants that were better recognized using TOMTOM^33^. For example, ACCTAA-4-TGTCNN and CCTAAC-3-TGTCNN together form an extended consensus ACCTAAC that significantly matches MYB TF binding sites (Fig. 1e). Still, individual biAuxRE appeared in several different alignments, i.e. ACCTAA-4-TGTCNN also aligns with CTAATC-2-TGTCNN or TACCTA-5-TGTCNN, suggesting they are part of larger or distinct constellations of motifs (Fig. 1e).

To test this hypothesis, we mapped the identified biAuxRE’s to the Arabidopsis genome (Supplementary Table 5). Indeed, this revealed many clusters of diverse biAuxRE compositions (henceforth “modules”) that contained more than one predicted bipartite element (located nearby, less than 15 nt, or partially overlapped). In total, 1136 promoters with regulatory modules were identified: most of them (1068) contained one module, 50 genes contained two, and 18 harboured three or more (Supplementary Table 6). There was a strong association between the presence of the predicted auxin-responsive module and auxin response in the transcriptomes for both activated and downregulated genes (p-value < 2.2E-16, Fisher’s exact test; Fig. 1f). Among the genes with multiple regulatory modules were members of the *SAUR*, *GH3*, and *Aux/IAA* families, which are rapidly and strongly activated by auxin^34^.

Finally, we analyzed how the regulatory modules distribute relative to experimentally determined transcription factor binding sites. For this, we used the Plant Cistrome DAP-Seq profile collection, which contains 568 peak sets for 387 TFs^18^. 66% of the predicted regulatory modules overlapped at least one TF peak; 13.6% of which overlapped with ARF-binding profiles (10.8% overlapped with ARF5 peaks and 2.9% with ARF2). There was a significant enrichment of those modules that overlap both ARF and another TF peak set compared to modules with only ARF peaks (p-value = 1.86E-09 for ARF5 and 1.4E-04 for ARF2, Fisher’s exact test). Although DAP-Seq is an *in vitro* method that can not account for TF-TF interactions, the analysis suggests that the predicted regulatory modules overlap the binding regions for different TFs.

### The presence of biAuxRE’s accurately predicts auxin-responsive transcription

The biAuxRE’s were identified based on their enrichment in genes that are auxin-regulated. However, it is unclear how predictive their presence in a given gene is for auxin-dependent expression. To test this, we selected 40 biAuxRE’s that have not previously been linked to auxin response, but showed the highest significance level in the permutation test. We randomly selected a gene possessing each one of the elements (Fig. 2a; Supplementary Table 7) and tested by qRT-PCR if the expression of these genes changed in roots, cotyledons, or seedlings after treatment with 1 μM of the synthetic auxin 2,4-D for one hour. For most of the selected genes, transcript level changed in response to auxin in at least one tissue (Fig. 2a). Following this large-scale but superficial testing, we selected ten up-regulated and ten down-regulated genes for further analysis. Arabidopsis seedlings were treated with 1 μM 2,4-D for 15, 30, 60, 180, or 360 minutes, after which qRT-PCR for these 20 genes was performed on roots. During this time-course auxin treatment, we examined how the transcriptional activity of genes with predicted biAuxRE’s in their promoters changed over time (up-or down-regulation; early or late response). Most of the chosen genes were dynamically up-or down-regulated in the early auxin response (Fig. 2b). The *AGP22*, *MYB40*, *BRL2*, and *ARP1* genes showed significant downregulation 2-3 hours after treatment, which is in accordance with the notion that their biAuxRE’s were associated with downregulated genes in the late response (Supplementary Tables 2, 5).

**Figure 2.**
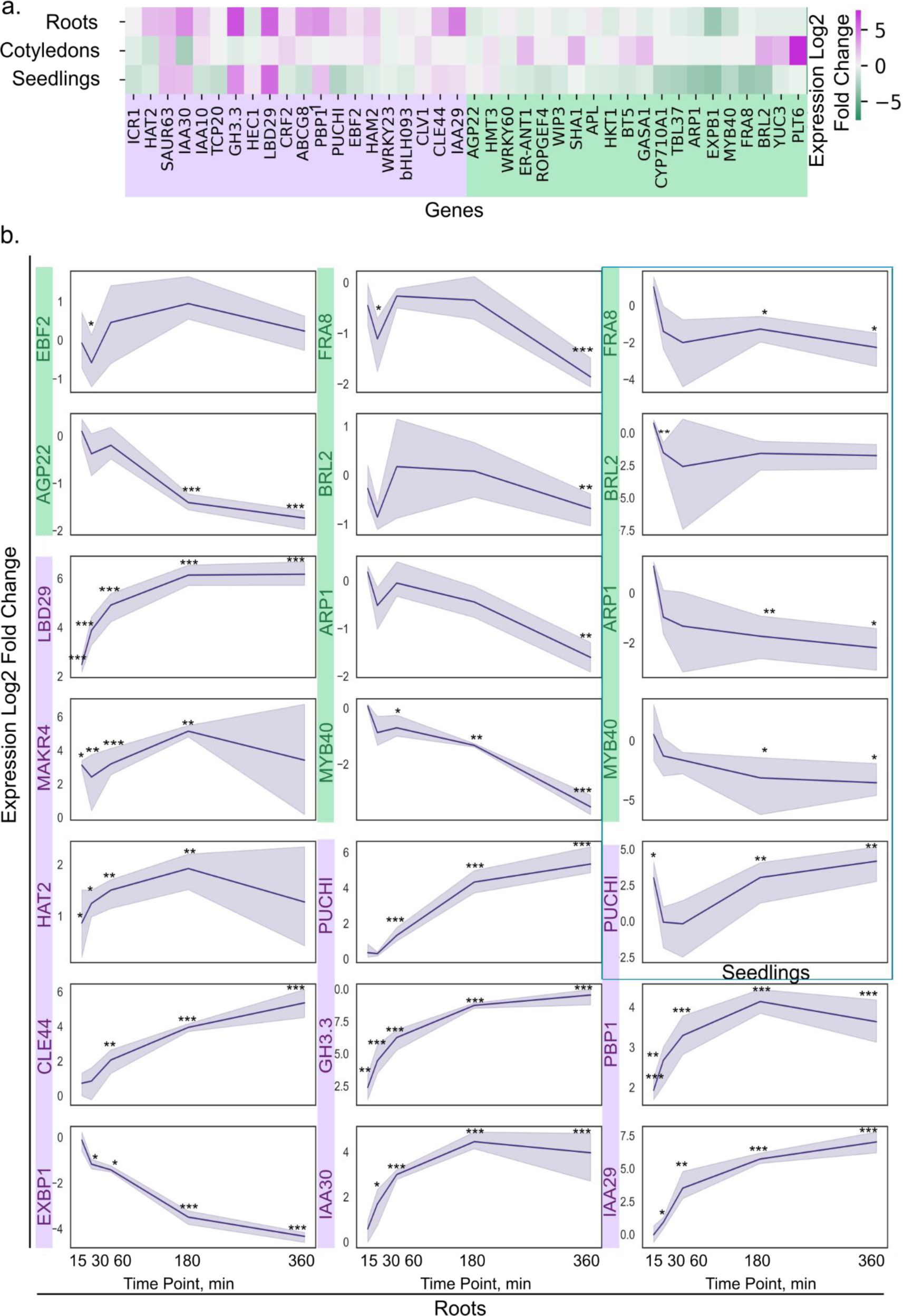
Genes with predicted bipartite auxin response elements show auxin-sensitive transcriptional response. **a**. A heatmap for gene expression changes in response to 1 hour of 1 μM 2,4-D treatment in roots, cotyledons, and whole seedlings (Supplementary table 7). **b**. Dynamic expression changes in response to 2,4-D treatment of 5 genes in roots and seedlings and for 10 genes in roots are shown with a separate graph for each gene (T-test p-value < 0.001 - ***, < 0.01 - **, < 0.05 - *).

We tested only seedling tissues at a single developmental stage, yet biAuxRE enrichment can reflect auxin control of any tissue or stage. We therefore consider the validation rate an underestimation of true values, and conclude that bipartite AuxRE’s accurately predict auxin responsiveness in Arabidopsis genes.

### Functionality of biAuxRE’s *in vivo*

To address the biological function of the predicted regulatory elements in the promoters, we mutated biAuxRE’s in the context of a full promoter by replacing either the TGTCNN or its coupling elements with TTTTTT. Each promoter was fused to a nuclear triple GFP (n3GFP). For potential ARF-binding repeats, we tested the DR1 element (TGTCNN-1-TGTCTC) in *LATERAL ORGAN BOUNDARIES-DOMAIN 29* (*LBD29; AT3G58190).* Furthermore, we tested the DR13 (ATGTCTN-13-TGTCNN) element in *MEMBRANE-ASSOCIATED KINASE REGULATOR (MAKR4; AT2G39370)* and the DR15 (TGTCTG-15bp-TGTCNN) element in *INDOLE-3-ACETIC ACID INDUCIBLE 30* (*IAA30; AT3G62100).* The *MAKR4* promoter additionally contained a TGTCNN-10-AATCCA biAuxRE that forms a module with DR13. The module associated with DR15 in *IAA30* was more complex, containing in addition a TGTCNN-10-GGTCAA and a CAAATA-4-TGTCNN element. We tested CAAATA-4-TGTCNN also in *GATA TRANSCRIPTION FACTOR 23* (*GATA23; AT5G26930*).

For each variant of upstream regulatory regions, we generated multiple homozygous lines and studied GFP expression in roots using confocal microscopy. Given the time needed for transcription, protein expression and folding, we analyzed roots after 3 to 9 hours of auxin or mock treatment, depending on the line. All the lines expressing n3GFP from intact promoters showed significant auxin responses under these conditions.

LBD29, MAKR4, and GATA23 mediate lateral root formation and their expression is induced by auxin^35–37^. Probably because of the dynamic nature of the lateral root initiation process, the gene expression patterns of *LBD29* and *GATA23* in the differentiation zone show great variability, especially upon auxin treatment, making it difficult to quantify the auxin-induced changes. To overcome this challenge, we quantified gene expression for *LBD29* in the root tip where it also expresses (Fig. 3e,f), while for *GATA23* we performed qRT-PCR for the *GFP* mRNA. *MAKR4* intact promoter expression was more focused, and we quantified the GFP intensity directly at the lateral root primordia sites (Fig. 3j-m).

**Figure 3.**
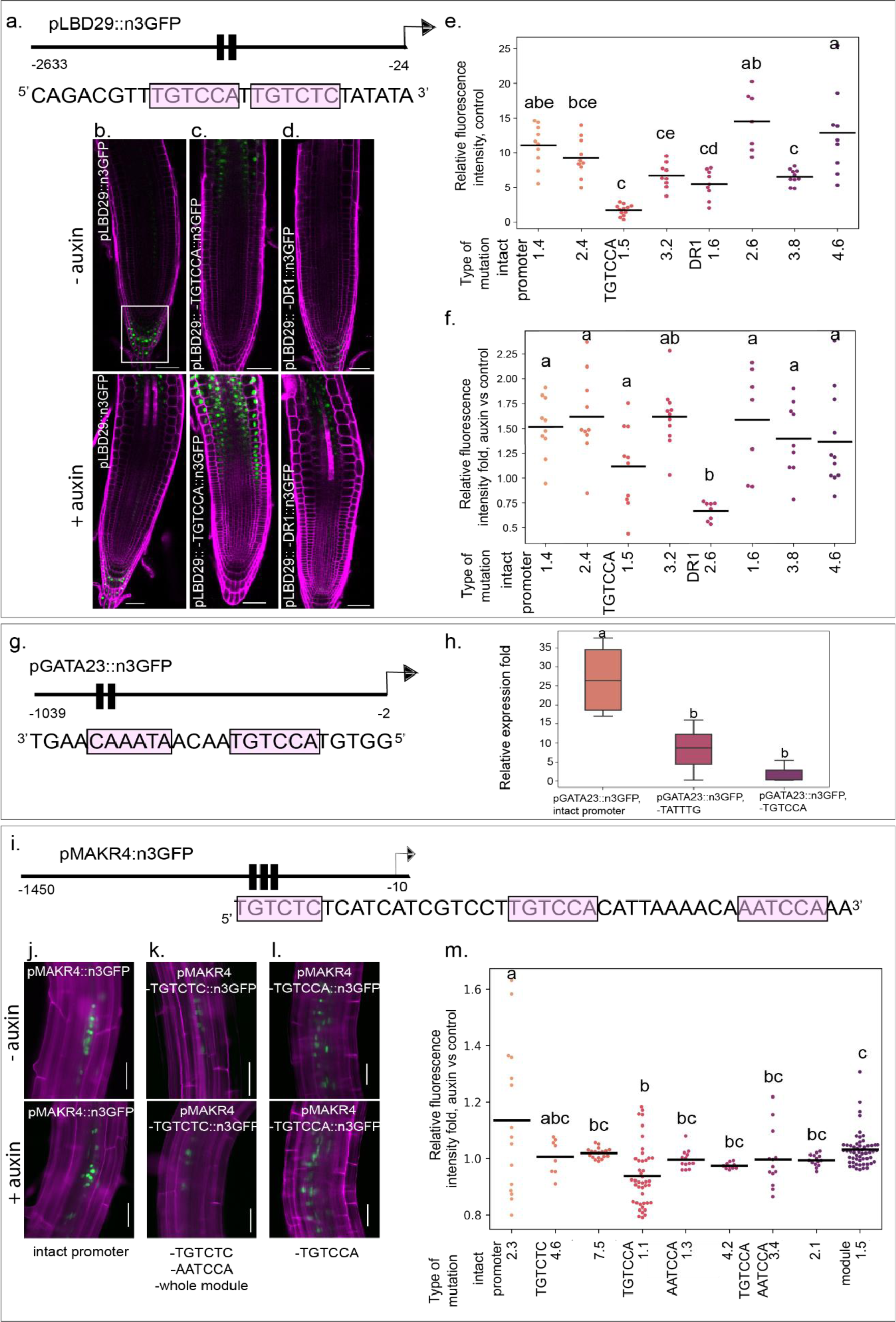
The predicted biAuxRE in *LBD29, GATA23, and MAKR4* upstream regions are required for gene expression in control and their responses to auxin. **a**. Scheme of the LBD29 upstream region with the predicted biAuxRE included in the analysis. **b-d**. Confocal images of reporter lines with intact and mutated promoter in control and after 7 hours of 1 µM 2,4-D treatment. The zone used in the GFP signal quantification is marked out with a white frame. Imaging was performed with a Leica SP8 microscope. Scale bar 50 μm. **e-f**. Results of GFP fluorescence intensity quantification in Arabidopsis root tips in LBD29 reporter lines in control **(e)** and after 7 hours of 2,4-D treatment versus control **(f)**. The relative fluorescence intensity is shown on the Y-axis and the lines used in the analysis on the X-axis. Statistical analysis: ANOVA, p-value < 0.05 with post-hoc Tukey’s test. **g**. Scheme of the GATA23 upstream region with predicted biAuxRE taken into analysis. **h**. qPCR results on the GFP expression in GATA23 reporter lines after 6 hours of 1 µM 2,4-D treatment (T-test p-value < 0.001 - ***, < 0.01 - **, < 0.05 - *). **i**. Scheme of the MAKR4 upstream region with the regulatory module included in the analysis. **j-l**. Representative images of MAKR4 patterns in the lateral root primordia under control conditions and after 4 hours of 1 µM 2,4-D treatment, showing changes when the regulatory module was mutated. Scale bar 100 μm. **m**. Fold change response to auxin in GFP fluorescence intensity quantification in early lateral root primordia in MAKR4 reporter lines. Imaging was performed using an LSM 780 NLO confocal microscope. Statistical analysis: ANOVA, p-value < 0.05, with a post-hoc Tukey’s test.

In the root tip, *LBD29* is expressed in the columella and weakly in the pericycle, and auxin treatment enhances this expression (Figure 3a,b). Disruption of one of the DR1 parts (TGTCCA) in the *LBD29* promoter led to a significant decrease in gene expression in these tissues in control conditions; however, the gene expression changes in response to auxin in this domain were not consistent between the individual lines. Notably, we observed higher auxin-induced expression in the proximal part of the meristem towards the transition zone (Fig. 3b-d). Unfortunately, our attempts to independently mutate the second DR1 motif were unsuccessful, but we managed to mutate both DR1 motifs together. Such lines showed expression level changes in the control, but the auxin response did not change relative to the intact promoter in three out of four lines (Fig. 3e,f). Although we see clear differences in the activities of intact promoters or their mutated versions, we can only conclude that both DR1 and TGTCCA mutations destabilize the expression pattern in control, and the TGTCCA mutation inverts the expression domain in the root tip upon auxin treatment. Taking into account that this genomic region has been shown to bind ARF7 in an electrophoretic mobility shift assay^35^, our results suggest that the DR1 element might provide for ARF7/19-mediated *LBD29* spatial patterning.

The intact *GATA23* promoter showed pronounced auxin responsiveness in the qPCR experiment with *GFP* mRNA, while mutation of either the AuxRE or the coupling element led to a strong decrease in auxin responsiveness (Fig. 3h). Either single, double, or triple site mutations in the predicted *MAKR4* module led to a loss of auxin response relative to control (Fig. 3j-l). However, we noticed a difference upon mutation of the central TGTCCA site in DR13; the gene expression domain was enlarged in this line, still showing no response to auxin (Fig. 3l). *GATA23* and *MAKR4* have not been reported to be direct ARF targets, the presence of biAuxRE in their promoters implies their expression is mediated via ARFs together with other TFs.

This analysis confirmed the functionality of non-standard repeats DR1, DR13, and the unannotated bipartite elements TGTCNN-10-AATCCA and CAAATA-4-TGTCNN. Although disruption of the elements in *GATA23* and *MAKR4* promoters reduced the genes’ expression in response to auxin, these elements were not essential for GATA23 and MAKR4 expression in control conditions. In contrast, the DR1 element played a more pronounced role in shaping the *LBD29* expression in control conditions. Interestingly, the specific effects of single and double site mutations in DR1 suggest that it might serve for TF competition rather than cooperation. Similarly, the central TGTCCA element in the *MAKR4* module might play a more specific role in narrowing down the MAKR4 expression domain.

### Analysis of the structural organisation of an auxin-response module in *IAA30*

The *IAA30* gene has a particular constellation of motifs, with three biAuxRE’s arranged in what resembles a module (Fig. 4a). In the reverse strand, central TGTCAT forms two biAuxRE’s with a WRKY-box TTGACC to the left and an A/T-rich motif TATTTG to the right.

**Figure 4.**
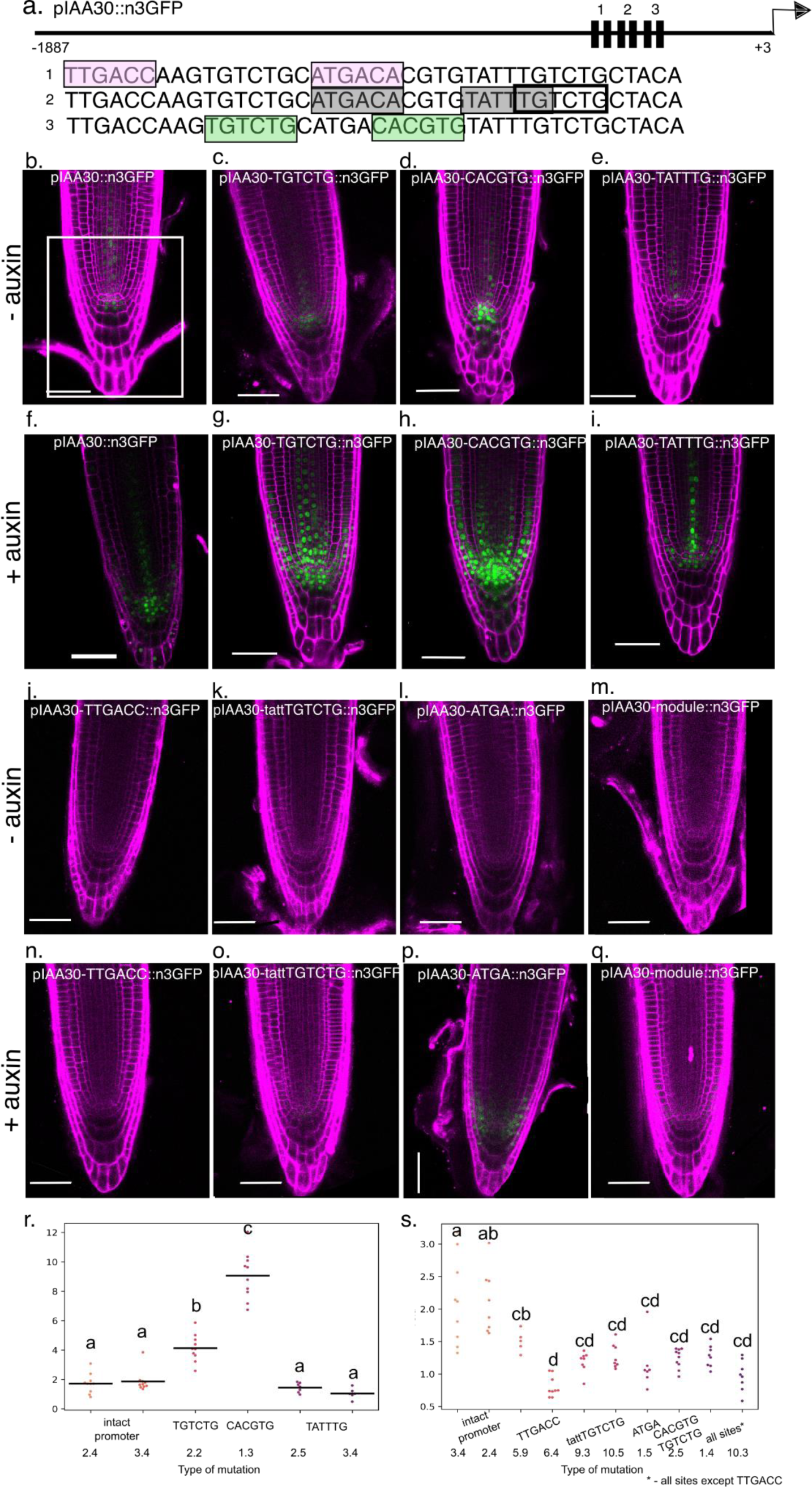
The predicted cis-regulatory module in the IAA30 upstream region is required for native auxin response. **a**. Scheme of the IAA30 regulatory module included in the analysis. **b-q**. Confocal images of reporter lines with intact and mutated promoter in the control and after 9 hours of 1 µM 2,4-D treatment. The zone used for quantification is marked with a white frame. Scale bar 50 μm. **r,s**. Results of GFP fluorescence intensity quantification in Arabidopsis root tips in IAA30 reporter lines. Fold change response in the relative fluorescence intensity is shown on the Y-axis, and the lines used in the analysis - on the X-axis. Statistical analysis: one-way ANOVA with post hoc Tukey’s test. **b-i.** Imaging was performed with Leica SP8. **j-q**. Imaging was performed with Zeiss Meta 780 NLO.

In the direct strand, a DR15 element partially overlaps the A/T-rich motif with one of the sites. Moreover, we found one more relevant motif in the module, a G-box CACGTG that partially overlaps the central TGTCAT site. CACGTG has not been found as a coupling motif in this study, but the composite AuxREs with CACGTG have been shown earlier in auxin-responsive genes^27,38^; and we found the single motif earlier as associated with transcriptional auxin response^21^. Thus, we proposed six motifs that make up the regulatory module and made eight promoter variants that changed one or more of these motifs to direct GFP expression.

The intact *IAA30* upstream regulatory region drives gene expression in the root stem cell niche; the gene expression is increased and the gene expression domain is enlarged upon auxin treatment (Fig. 4b,f). Disruption of all sites in the module abolished gene expression in control and its auxin responsiveness (Fig. 4m,q), the same effect was observed in the single site mutations for the starting WRKY-box and the closing taattTGTCTG motif, the right part of DR15 (Fig. 4i,n,k,o). Intriguingly, mutations in either the left part of DR15, TGTCTG, or CACGTG led to increased expression with a broader expression pattern both in control and upon auxin treatment (Fig. 4c,g,d,h). Notably, mutations of both sequences together (TGTCTG and CACGTG) caused a loss of auxin responsiveness (Fig. 4s). Since CACGTG overlaps with the central TGTCAT element (Fig. 4a), the double mutation disrupts all the predicted biAuxRE destroying the regulatory module’s integrity, which explains the opposite effects of single and double mutations. Also, when we mutated only the central TGTCAT element, keeping CACGTG untouched, this significantly decreased the gene expression and its response to auxin (Fig. 4l,p); so the overlapping motifs play opposite activating and inhibitory roles in auxin response. Another two overlapping motifs CAAATA/TATTTG and tattTGTCTG (part of DR15) show the same activating effects, but the role of tattTGTCTG might be higher (Fig. 4e,i,k,o).

In summary, five motifs (TTGACC, TGTCTG, TGTCAT, CACGTG, and tattTGTCTG) appear to be functional in the studied *IAA30* promoter region, and we therefore conclude that this indeed acts as a regulatory module. The presence of all of the motifs seemed to be required for gene expression in the root stem cell niche and its response to auxin. However, the relationships between the elements in the module were not trivial, with DR15 showing the opposite effect when one or another part is mutated and with a functional G-box that inhibits the regulatory module activity. This is evidence of the complexity of the binding sites as part of regulatory modules and showed the necessity to study not only single and bipartite elements but also bipartite elements as a part of regulatory modules.

### Identification of TFs binding the *IAA30* regulatory module

Since the cis-regulatory logic of the auxin-mediated mechanism of *IAA30* transcriptional regulation, as inferred from mutations, appears complicated, we decided to address the mechanism underlying its regulation by TFs. A Yeast One Hybrid (Y1H) screen was performed to find candidate regulators of *IAA30*. We screened the interaction of 1957 TFs^39^ (Fig. 5a) with a triple repeat of the *IAA30* regulatory module. In this nearly saturating screen, 24 positive interactions were found (Fig. 5b; Supplementary table 8). Only about half of the ARF TFs are represented in the library, and we did not recover any of these on the screen. Given that we used the entire regulatory module as a screening bait, we next mutated the CACGTG and TATTTG motifs to identify TFs that likely bind to these two elements or overlapping the TGTCNN motifs. Twenty TFs (bZIP (AT5G42910), bZIP3, bZIP16, bZIP42, bZIP44, bZIP48, LBD18, LBD20, LBD30, ABF2, ABF4, GBF1, GBF2, GBF3, ABI5, WOX3, MADS (AT5G27810), WRKY7, C2H2, HEC2) lost binding in this assay when the CACGTG element was mutated (Fig. 5c). The remaining set of TFs (LBD19, LBD31, bZIP12, ABF3) did not require the presence of the CACGTG or tattTGTCTG elements (Fig. 5a,b), and likely bound to TTGACC or TGTCNN, or to other unknown binding sites in this regulatory module. When compared with TF’s DAP-Seq profiles^18^, the *IAA30* regulatory module was found to overlap multiple bZIPs and WRKY binding regions, as well as the ones for ARF5 and ARF2 (Supplementary Figure 1).

**Figure 5.**
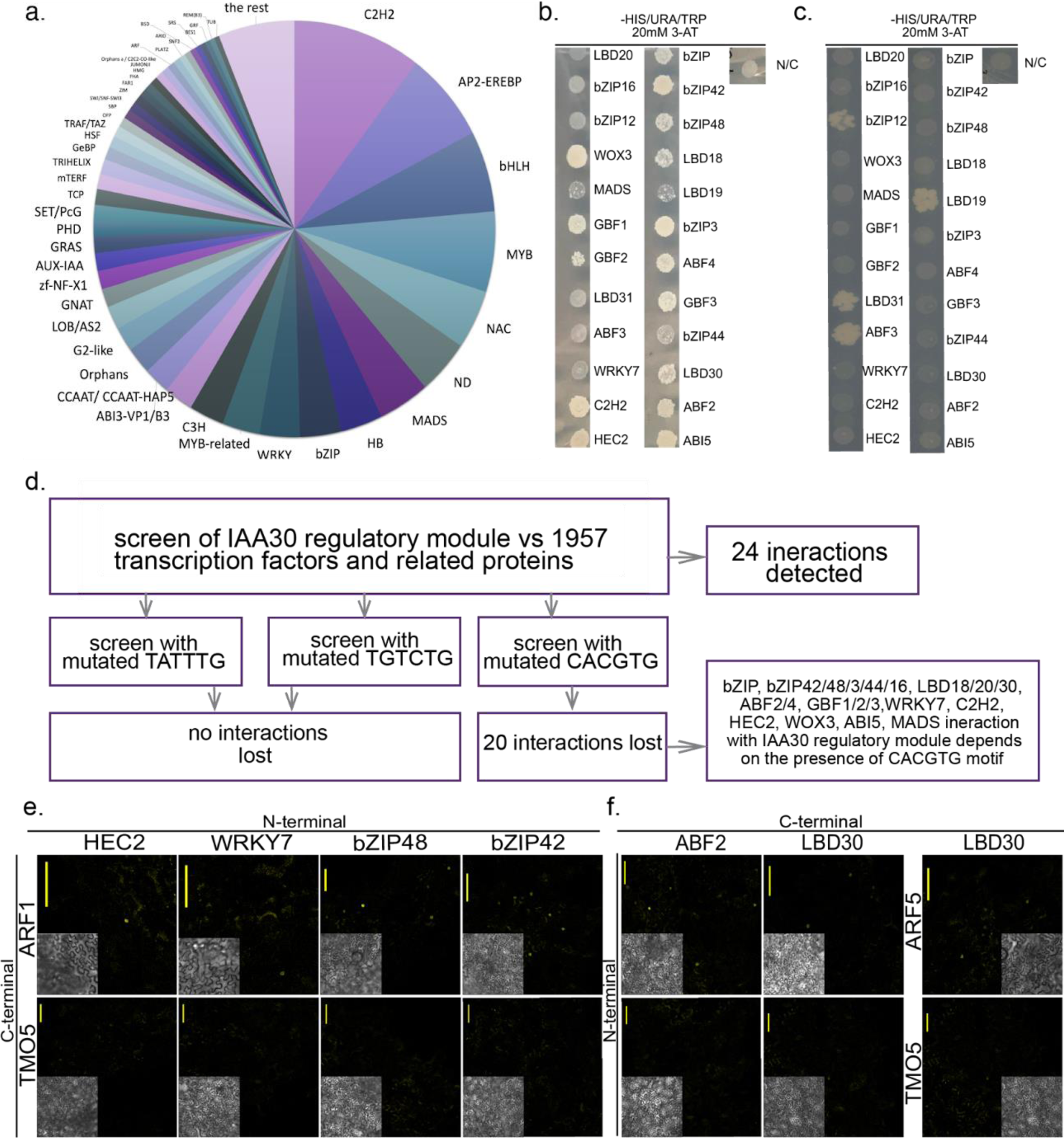
Identified DNA-protein and protein-protein interactions with the IAA30 regulatory module. **a**. Transcription factors library used for yeast-one-hybrid (Y1H) assay^39^. **b**. Transcription factor interactions with the *IAA30* regulatory module identified via Y1H. D. Interactions with the *IAA30* regulatory module with the mutated CACGTG site. **d**. Schematic summary of the Y1H results. **e-f**. Revealed interactions of ARF1 and ARF5 with potential *IAA30* regulators in BiFC assay. Scale bar 100 μm. **e**. Interactions of C-terminal YFP part fused with ARF1/5 and the N-terminal YFP part fused with transcriptional factors detected in the Y1H assay. **f.** Interactions of the N-terminal YFP part fused with ARF1/5 with the C-terminal YFP part fused with transcriptional factors detected in the Y1H assay.

The dependence of auxin-mediated *IAA30* expression on the identified biAuxRE could stem from protein-protein interactions between the TFs binding these elements and the ARFs. We therefore experimentally tested interactions between proteins recovered in the Y1H screen and the A-class ARF ARF5 and the B-class ARF ARF1, using a bimolecular fluorescence complementation assay (BiFC, split-YFP). For each TF, we generated an N-terminal fusion with the N-terminal (Nt) part of YFP, and an N-terminal or C-terminal fusion of the C-terminal (Ct) part of YFP. When co-expressed in *Nicotiana benthamiana* leaves, we detected interactions between LBD30 and ARF5 and between bZIP42, bZIP48, ABF2, WRKY7, HEC2 and ARF1 (Fig. 3e,f). Thus, the TFs identified in this study are potential interaction partners of ARFs in the auxin-mediated transcription of *IAA30* and other genes.

If any of these TFs play a role in the control of *IAA30* expression, one would expect some degree of (anti-)correlation in their cellular expression patterns. We tested this by comparing the *IAA30* pattern with that of all TFs recovered in the yeast screen and BiFC assay using the RootCellAtlas^40^ which integrates single-cell transcriptomics data from multiple scRNA-Seq datasets. We found that *ABF2*, *WRKY7*, and *bZIP42* have expression domains partially overlapping with either *ARF1*, *ARF5*, or *IAA30* (Fig. 6a). Several other TFs did not show detectable expression in this root domain, and were not considered further.

**Figure 6.**
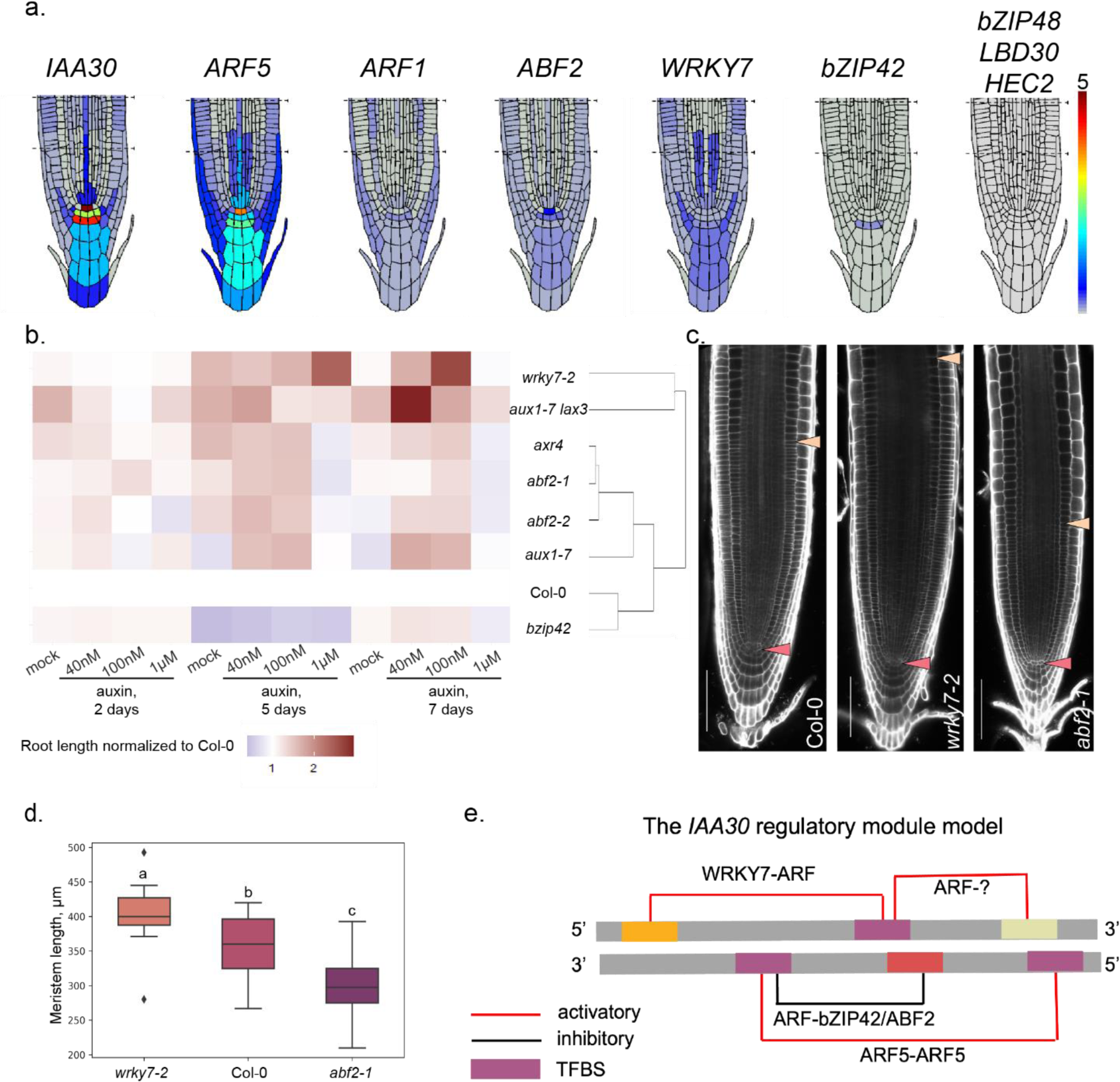
The functional model for the IAA30 regulatory module. **a**. Expression patterns of IAA30 and the identified TF regulators in the root tip. The data was taken from the RootCellAtlas web-service^40^. **b**. Primary root length differences in control (Col-0), auxin-related mutants *(axr4, aux1-7, aux1-7 lax3),* and the candidate gene mutants *(wrky7-2, bzip42,* and *abf2-1, abf2-2)* during auxin treatment course. Plants were transferred at 3 DAG to 1/2MS, 40 nM, 100 nM, or 1 µM IAA with three consecutive measurements of the root length (2, 5, and 7 days after transfer). The root lengths were normalized to the Col-0 data and the integrated data was subjected to hierarchical clustering. The detailed statistical data is in Supplementary Figure 3. **c**. Root meristem anatomy in the respective mutant lines (5 DAG, control). QC is marked by red arrowhead, the meristem border by yellow arrowhead. Scale bar 100µm. **d**. Meristem length (µm) of the mutant lines and Col-0 reflected on **c**. (T-test p-value < 0.01 - **, < 0.05 - *). **e**. The model of IAA30 regulatory module functioning. Individual sites are denoted with blocks of different colours, and validated bipartite elements are shown with black (inhibitory) and red (activatory) lines. Each biAuxRE has an assigned pair of transcription factors’ partners.

None of the ABF2, WRKY7 or bZIP42 TFs have been reported to control meristem development or auxin response. We therefore obtained insertion mutants in each and characterized phenotypes in root development and auxin response. For comparison, we included well-known auxin response mutants *axr4, aux1-7 and aux1-7 lax3*^41–43^. Responses of primary root growth to different auxin concentrations (0 nM, 40 nM, 100 nM, or 1 µM IAA) were compared between the candidate gene mutants, wild-type control (Col-0) and auxin-related T-DNA insertion mutants (Fig. 6b, Supplementary Figures 2, 3). While the auxin-responsive growth pattern of the primary root in the *bzip42* mutant largely mimicked the one in wild-type, *wrky7-2* and *abf2* alleles showed significantly different responses. Auxin-responsive growth in *abf2-1* and *abf2-2* was similar to that in *axr4* and *aux1-7,* while *wrky7-2* showed response patterns resembling *aux1-7 lax3* mutant. The main difference between those two groups of mutants was in the reaction to high (1 µM) IAA concentrations, which was inhibitory for *abf2* group and stimulatory for the *wrky7* group. As auxin plays a major role in root meristem patterning, we also measured the meristem length in control and mutant genetic background. While the root meristem anatomy was unchanged in *bzip42,* it was reduced in *abf2-1* and extended in *wrky7-2* mutants (Fig. 6c-d).

Altogether, the data on motif mutagenesis, DNA-protein and protein-protein interactions, and root phenotypes allow us to propose a structural-functional model for the *IAA30* regulatory module (Fig. 6f). The model suggests co-regulation of *IAA30* transcriptional activation via WRKY7-ARF and ARF5-ARF5 dimers that bind non-overlapping motifs within the regulatory module. *IAA30* transcription is attenuated via the ABF2-ARF (Supplementary Figure 1) complex and the bipartite element that overlaps with both activatory biAuxRE. Examining the expression levels of *IAA30* and its regulators under control conditions (Fig. 6a) suggests that the ARF5 activator exerts the most significant influence, while other TFs modulate its expression range. We hypothesize that the co-regulators may assume a more prominent role in response to fluctuations in auxin levels under changed environmental conditions. To investigate this supposition thoroughly, a comprehensive analysis of the expression patterns of all implicated factors under various conditions is necessary.

## Discussion

The prevailing perspective on transcriptional regulation mechanisms suggests that transcriptional enhancers or regulatory modules harbour TF binding motifs arranged in specific configurations, also known as syntax^44,45^. TF binding site orientation and spacing can crucially influence the affinity of TF complex binding to DNA^16,17^. Based on this syntax, TFs bind the motifs either cooperatively or competitively, subsequently facilitating the recruitment of co-activators or co-repressors that regulate gene activity. The identification of regulatory modules typically involves DNA-protein binding assays, including DAP-Seq, ChIP-Seq, or ATAC-Seq. Current methodologies provide TF binding sites and syntax inference through bioinformatics or machine-learning analyses of the regulatory modules, in a top-down approach.

In this study, we adopted an alternative bottom-up approach to uncover functional regulatory modules, treating them as arrangements of motifs that are consistently overrepresented in the promoters of the genes responsive to auxin. In an unbiased search without predefined assumptions for ARF partners, we screened through all possible hexamers pairing with the potential ARF binding site TGTCNN, identifying those pairs that exhibited consistent overrepresentation within the promoters of auxin-mediated genes (Fig. 1a). With a similar approach, we had showcased the systematic presence of single cis-regulatory motifs^21^ and TGTCNN repeats^16^ in auxin-sensitive promoters. Next, the genome-wide mapping of the identified biAuxRE highlighted potential regulatory modules. Although the auxin-responsive modules were identified purely bioinformatically, 66% of them overlapped at least one DAP-Seq peak from Plant Cistrome collection^18^; moreover, there was a strong association between the presence of the predicted module and auxin activation or repression (Fig. 1f). Many primary auxin-responsive genes, like *SAUR*, *GH3*, and *Aux/IAA* genes, possess more than one module in their promoters (Supplementary Tables 5, 6).

The classical auxin transcriptional response encompasses ARF-binding repeats anchoring ARF dimers. A few types of TGTCNN repeats have been identified and experimentally validated: DR5, IR8, and IR13 ^16,17,19,32^. Notably, in this study, we detected all of them as significantly associated with auxin-responsive transcriptional activation (Fig. 1c). Moreover, we detected new yet unknown repeats overrepresented not only in promoters of genes in early auxin activatory responses but also in late activatory and inhibitory responses (Fig. 1c, Supplementary Table 3). Here, we experimentally validated three such repeats: DR1 in *LBD29*, DR13 in *MAKR4*, and DR15 in *IAA30* promoters. Unlike other validated repeats, mutations of individual motifs within DR1, DR13, and DR15 lead to contrasting effects. The experimental results for DR1 and DR15 suggest a competitive rather than cooperative nature of TF binding to the repeats in *LBD29* and *IAA30* promoters (Figure 3a-f; 4c,g); and in *MAKR4*, one of the DR13 half-sites likely has an additional, non-cooperative role to narrow down the expression domain (Figure 3i-m). This data reveals new dimensions in the exploration of classical transcriptional auxin responses, unveiling the hidden intricacies of ARF-binding repeat-mediated responses.

Still, existing knowledge on the auxin signaling pathway is not sufficient to explain the complexity and diversity of processes that are regulated by auxin locally^1,7^. Our hypothesis posited the involvement of non-ARF TFs in local auxin responses, working in conjunction with ARFs to govern diverse developmental processes. Among the predicted coupling elements, binding sites for several TF families were more abundant: bZIP, MYB/MYC, WRKY, TCP, C2H2, and AP2/ERF (Fig. 1d). Using the predicted *IAA30* regulatory module to identify ARF partners, we identified 24 TFs that bind this region in Y1H assay; for 20 of these, DNA binding depended on the central CACGTG motif, or its overlapping motif TGTCAT (Figure 5b-d). The spectrum of upstream regulators included the members of LBD, bZIP (including ABFs and GBFs), bHLH, MADS, and WOX families. Previous research has highlighted the involvement of bZIP TFs as chromatin remodeling co-factors in auxin regulation^46^. Furthermore, we previously found an interaction between GBF1 and GBF2 with G-box and ARF5 in the context of vasculature development^28^. LBD factors have been identified before as mediating auxin-mediated processes^35,47^. For instance, *LBD18* is activated by auxin through ARF7/19 in lateral root development, moreover, it dimerizes with ARF7/ARF19 and activates *ARF7* transcription in a positive feedback loop^48^.

To further refine the list of upstream regulators, we tested in BiFC assay which of them form heterodimers with ARF1 and ARF5. The resulting list included bZIP factors ABF2, bZIP42, and bZIP48; WRKY7, LBD30 and HEC2 (Figure 5e,f). The functions of bZIP42 and bZIP48 in plant development remain largely unexplored; here we also have not detected their roles in the root meristem (Fig. 6a-b). Similarly, HEC2 was found to be absent in the root, with its primary role being associated with the regulation of female reproductive tract development^49^. Notably, a close homolog of HEC2, HEC1, has been tightly integrated into the auxin-signaling network during flower development^50^. It is a question for further studies if detected ARF1-HEC2 interactions are operational during flower development.

LBD30, also known as JLO, controls embryonic patterning together with ARF5^51^. The novel data revealing protein-protein LBD30-ARF5 and DNA-protein *IAA30*-LBD30 interactions (Fig. 5) offers fresh insight into the functional role of LBD30/JLO within the LBD30-ARF5-IAA30 regulatory module.

Here we discovered the role of WRKY7 and ABF2 in mediating auxin response in the root. Pathogen-induced WRKY7 has been implicated in the negative regulation of basal resistance to *Pseudomonas syringae*^52^ and is known to mediate the unfolded protein response^53^. Furthermore, ABA-responsive ABF2, also known as AREB1, has been found to facilitate physiological adaptation to drought stress^54^, to mediate salt stress^55^, and to elicit a response to nitrate^56^. Neither of these TFs has been previously associated with the regulation of the auxin response or meristem activity. Our findings demonstrate that both WRKY7 and ABF2 can bind with ARF proteins and bind to the *IAA30* promoter (Figure 5). Moreover, both mutants displayed reduced sensitivity of primary root growth to auxin (Supplementary Figure 3), resembling the behaviour observed in other auxin-insensitive mutants such as *aux1-7, axr4*, or *aux1-7 lax3* (Figure 6b). The mutants responded differently to high auxin concentrations, though, with *wrky7-2* and *aux1-7 lax3* mutants being less sensitive, and the rest being more sensitive than Col-0. Another difference was detected for the meristem length, although *abf2* mutants exhibited shorter meristem lengths compared to the Col-0 plants, the meristem of *wrky7-2* mutant was longer (Fig. 6c-d). Auxin has been identified as a crucial factor in integrating environmental signals into plant root development^57^. Through our discoveries regarding WRKY7 and ABF2, we have shed light on the concealed layer of transcriptional auxin response co-regulation, being potentially influenced by environmental cues through the identified biAuxRE’s and their modules. To comprehensively unravel this co-regulation network, a systematic analysis of composite responses to auxin under various stresses will be imperative.

The transcriptional machinery underlying auxin response is far more complex than the basic auxin signal transduction mechanism. According to the predictions, dozens of TFs are involved in co-regulation of auxin-mediated processes. Further investigations are required to address the association of bipartite elements with particular TFs, processes, tissues and organs. That would allow us to construct a more complete network of auxin-mediated transcription regulation.

## Methods

### Plant material, growth conditions and treatments

Plants were grown at 22℃ under long daylight mode (16h light, 8h dark). Seeds were surface sterilized and underwent 2-7 days of stratification at 4℃. Arabidopsis ecotype *Columbia-0* and *pGATA23::3nGFP* Arabidopsis plants for qPCR analysis were grown vertically on ½ MS for 5 days. 3 DAG plants were treated with 1μM 2,4-D on solid ½ MS. For antibiotics selection of T1, T2 and T3 of transcriptional reporter lines seedlings were initially grown on ½ MS with 50 mg/L kanamycin for 7 days. Then transformants were transferred to soil for further selection and propagation. *Nicotiana benthamiana* 3-week-old plants grown under standard conditions were used for BiFC experiments. Following lines were used in the root growth assay: Col-0, auxin-related mutants (*axr4*^41^, *aux1-7*^42^, *aux1-7 lax3*^43^), and T-DNA insertion lines *wrky7-1* (SALK_093993C) *wrky7-2* (SALK_093993), *bzip42* (SALK_047165C), and *abf2-1* (SALK_002984C), *abf2-2* (SALK_138855C), *lbd30* (SALK_134715), *bzip48* (SALK_110507). T-DNA insertion lines were verified by genotyping (Supplementary Table 9). 3 DAG plants were transferred on ½ MS media and 40 nM, 100 nM, and 1 µM IAA supplemented ½ MS media. *abf2-1, abf2-2, lbd30, and bzip48* lines were still segregating in the root length measurement experiments.

### Expression analysis

Plant material (seedling, roots or cotyledons) of 3 DAG Arabidopsis seedlings was flash frozen in liquid nitrogen and powdered with TissueLyser machine (Qiagen). RNA was isolated with TRIzol (Invitrogen) and RNAeasy kit (Qiagen). cDNA was synthesized from 1μg total RNA with iScript cDNA Synthesis kit (Biorad). Real-time PCR was measured on CFX384 RT-PCR detection system using iQ SYBR Green Supermix (Biorad) or its analogue EvaGreen (Sintol). Each reaction was performed in three technical and three biological replicates. *ACTIN2* was used as the reference gene. All primers used for qPCR analysis are listed in Supplementary Table 9.

### Generation of transcriptional reporters

All primers used for cloning are presented in Supplementary Table 9. To replace predicted motifs in promoters with the TTTTTT sequence we used the overlap extension PCR method^58^. Amplified with Fusion polymerase (Thermo Scientific) promoters’ versions were introduced into the *pPLV04_v2*^59^ backbone with SLICE cloning^60^ and confirmed with sequencing. *A. thaliana* plants were transformed with these constructs using the floral dip method^59^.

### Microscopy analysis

Confocal imaging of the reporter lines was performed on Leica SP8 system and on Carl Zeiss LSM 780 NLO. Roots were counterstained with 10 μg/mL propidium iodide (PI). GFP fluorescence intensity was quantified using ImageJ^61^ with the Analyze/Measure function. BiFC imaging was performed on Leica SP5 II system equipped with Hybrid Detectors. T-DNA insertion 5DAG lines were stained with PI and imaged on LEICA Stellaris confocal microscope.

### Yeast one hybrid screen

Triple repeat of *IAA30* regulatory module was cloned with Gateway Cloning kit (Life Technologies) in pMW2 and pMW3 backbones (Addgene plasmid #13350; http://n2t.net/addgene:13350; RRID: Addgene_13350) and transformed in PJ69-4α (*MATα, ade2, trp1-109, leu2-3, 112, ura3-52, his3-200, gal4, gal80, GAL2::ADE2, LYS2::GAL1:HIS3,met2::GAL7:lacZ*) (James et al., 1996). Transcription factors library with yeast PJ69-4A (*MATa trp-901 leu2-3, 112 ura3-52 his3-200 gal4D gal80D GALADE2 LYS::GAL1-HIS3 met::GAL7-lacZ*)^62^ carrying 1975 transcription factors cloned in pDEST22 backbone was used for the screen^39^. Yeasts were transformed following standard protocol^63^ and selected by testing autoactivation. Selection media without histidine, uracil and tryptophan with 3-amino-1,2,4-triazol (Invitrogen) added was used in the interactions’ screen. Detected interactions from the library (Supplementary Table 8) were confirmed with sequencing.

### Bimolecular fluorescence complementation (BiFC; Split-YFP)

CDS of transcription factors were cloned in the modified *pPLV26*^59^ backbone with C-terminal and N-terminal YFP parts. These plasmids were transformed in *Agrobacterium tumefaciens* used in the infiltration of *Nicotiana benthamiana* leaves. Two days after the infiltration leaf sections were cut and studied with confocal microscopy. TMO5 (AT3G25710, TARGET OF MONOPTEROS 5) and LHW (AT2G27230, LONESOME HIGHWAY) interaction was used as a positive control. TMO5 interaction with the testing TFs was used as a negative control.

### Bioinformatic analysis

We collected Arabidopsis gene promoters [-1500nt; TSS] from the TAIR10 genome version in the TAIR database^64^. To identify overrepresented biAuxRE’s, we applied the *metaRE* package^31^ to differentially expressed gene sets taken from multiple auxin-induced transcriptomes (Supplementary Table 1). The promoters for auxin early- and late-responsive, and for up- and down-regulated genes were analysed separately. The biAuxRE’s had the following structures: TGTCNN-spacer-NNNNNN or NNNNNN-spacer-TGTCNN on one of the DNA strands, where N could be any nucleotide (A, T, G, C), and the spacer length was set from 0 to 15 bp. First, we selected biAuxRE’s enriched in the promoters of differentially expressed genes over multiple transcriptomics datasets (Fisher’s method; meta-p-value<1E-5). Next we tested the robustness of biAuxRE’s overrepresentation with a permutation test, selecting the final biAuxRE’s set under permut-p-value <0.05 (Supplementary Table 2). TOMTOM tool^33^ (http://meme-suite.org) was used to annotate the coupling motifs to known TF binding sites (Supplementary Table 4). The regulatory modules (Supplementary Table 6) were identified as groups of predicted biAuxRE mapped to the promoters of the genes in a close vicinity (less than 15 nt distance, or partially overlapped with minimum 6 nt). To determine the association between the presence of predicted auxin-responsive modules and auxin response in the transcriptomes we compiled two 2×2 contingency tables: one for upregulated and one for downregulated auxin-responsive genes. These tables included counts of genes whose promoters either overlapped or did not overlap with the predicted modules. Fisher’s exact test was then applied to estimate the enrichment of these modules in the promoters of upregulated and down-regulated auxin-responsive genes relative to non-responsive genes.

We used the Plant Cistrome TF binding profiles collection^18^, which consists of 568 peak sets for 387 TFs, to assess the distribution of predicted regulatory modules relative to TF binding peaks. For genes with predicted regulatory modules, we mapped the DAP-seq peaks of each TF to the gene promoters and compiled two 2×2 tables for the presence/absence of the ARF5/ARF2 binding peak and the presence/absence of the peak for any other TF. Then we counted overlapping ARF-TF peaks on the predicted regulatory modules and evaluated overrepresentation with Fisher’s exact test.

## Author Contributions

VM and DW conceived and designed the study. DW and DN designed the experimental validation of biAuxRE’s. DN performed most of the experiments with input from AK and CA. VM, DN, and VVL conducted the bioinformatic analysis of biAuxRE’s. NO, DN, and VM analysed the bioinformatic output. DN, VM and DW wrote the manuscript with the input from other co-authors.

## Acknowledgments

Authors are grateful to the members of the sector of systems biology of plant morphogenesis (IC&G, RU), Department of Plant and Animal Biology (RU, NL) and Laboratory of Biochemistry (WUR, NL) for technical support and valuable feedback on the project. We especially thank Alexey Kochetov (IC&G, RU), Elena Zemlyanskaya (IC&G, RU), Victor Levitsky (IC&G, RU), Elmar van der Wijk (RU, NL), Tatyana Radoeva (WUR, NL) and Kuan-Ju Lu (WUR, NL) for their support and valuable comments. We thank the Center for Microscopy of Biological Objects (IC&G, SB RAS) and its head Sergey Baiborodin for excellent assistance with microscopic studies. DN thanks for technical support Alexandra Ceban, Pauline Clément, Nadine Eliasson, Anaïs Hautefeuille, Perrine Kergoat, and Elisa Schmid. VM is supported in part by Netherlands Organization for Scientific Research (NWO) OCENW.M20.197. NO is supported by RSF no. 20-14-00140. VVL is supported by the budget project FWNR-2022-0020. The project was supported by Wageningen Graduate Schools (WGS) Sandwich PhD scholarship for DN.

